# Human resting-state electrophysiological networks in the alpha frequency band: Evidence from magnetoencephalographic source imaging

**DOI:** 10.1101/142091

**Authors:** R. Hindriks, C. Micheli, D. Mantini, G. Deco

## Abstract

In the resting-state, extended regions of the human cortex engage in electrical oscillations within the alpha-frequency band (7–14 Hz) that can be measured outside the head by magnetoencephalography (MEG). Given the accumulating evidence that alpha oscillations play a fundamental role in attentional processing and working memory, it becomes increasingly important to characterize their cortical organization. Event-related studies have demonstrated that attentional allocation can modulate alpha power selectively within the visual, auditory, and somatosensory cortices, as well as in higher-level regions. Such studies demonstrate the existence of multiple generators by exploiting experimental contrasts and trial-averaging. The identification of alpha generators from resting-state data alone has proven much harder and, consequently, relatively little is known about their organization: Apart from the classical visual, somatosensory, and auditory rhythms, it is unclear how many more generators can be observed with MEG and how they are organized into functional networks. Such knowledge, however, possibly enables to delineate separate cognitive, perceptual, and motor processes that co-occur in the resting-state and is therefore important. In this study we use the resting-state MEG data-set provided by the Human Connectome Project to identify cortical alpha generators and to characterize their organization into functional networks. The large number of subjects (*N* = 94), multiple scans per subject, and state-of-the-art surface-based cortical registration enable a detailed characterization of alpha in human cortex. By applying non-negative matrix factorization to source-projected power fluctuations, we identify 16 reliable cortical generators in each hemisphere. These include the classical sensory alpha rhythms as well as several additional ones in the lateral occipital and temporal lobes and in inferior parietal cortex. We show that the generators are coordinated across hemispheres and hence form resting-state networks (RSNs), two of which are the default mode network (DMN) and the ventral attention network (VAN). Our study hence provides a further subdivision of RSNs within the alpha frequency band and shows that these RSNs are supported by alpha generators. As such, it links the classical literature on human alpha with more recent research into electrophysiological RNSs.

## 1 Introduction

Human alpha (7–14 Hz) oscillations were first observed using scalp electroencephalography (EEG) almost a century ago [5] and their physiological origin was confirmed in the late sixties using magnetoencephalography (MEG) [12]. It was not until about a decade ago that their involvement in selective attention has been firmly established. Alpha oscillations reflect fluctuations in cortical excitability [61] through deployment of selective attention [36, 34, 33]. For instance, allocating attention to one hemifield, leads to a decrease in alpha power in the contralateral visual cortex and to enhancement in the ipsilateral visual cortex [35]. This modulatory role of alpha has been established in the visual [74, 35, 60, 66], somatosensory [15, 24], and auditory [4, 70, 54, 71, 28, 73] domains as well as across domains [19, 67] and has been shown to have perceptual (and hence, behavioral) consequences. In these and related studies, alpha power is modulated in an repeatable way by controlled manipulation of selective attention. An interesting question that cannot be addressed by such experimental designs is how alpha is organized in the absence of such controlled manipulation, that is, during the resting-state. More specifically, which cortical regions spontaneously generate alpha and how are these generators coordinated? In controlled experiments such as those cited above, the existence of alpha generators is inferred by contrasting conditions and trial-averaging. For instance, in a spatially cued visual detection task, the existence of two independent generators within the visual system (dorsal and ventral) was inferred in such a way [9]. The existence of alpha generators in the resting brain cannot be inferred by using these techniques and this makes the above question particularly challenging to address.

The preclusion of using experimental contrasts and trial-averaging is one of the reasons that relatively little is known about the organization of spontaneous alpha in human cortex. Early EEG and MEG dipole studies have demonstrated that visual and primary somatosensory cortices spontaneously generate alpha oscillations. These oscillations have become known as the *visual alpha rhythm* and the *somatosensory mu rhythm*, respectively [27, 10, 46]. Although referred to as *rhythms*, visual and somatosensory alpha constitute a multitude of independent oscillatory generators. This is inferred from the observation that MEG dipoles typically form two clusters; one located within the Calcarine sulcus and another within the parieto-occipital fissure [62, 27, 10, 46, 42, 22]. A difference between visual alpha and somatosensory mu seems to be that the latter is confined to the hand area of the primary somatosensory cortex [27], while visual alpha is more widespread. The reason and functional relevance for this difference is unknown. Besides visual alpha and somatosensory mu, there have been reports of a more lateral rhythm within the somatosensory cortex, originating from the upper lip of the Sylvian fissure [64, 56, 42], where the secondary somatosensory cortex is located. This rhythm, which is referred to as the *sigma rhythm*, is not found in all subjects, presumably due to signal cancellation within the parietal operculum, but the available evidence is inconclusive. The organization of spontaneous alpha in the temporal lobe is incompletely understood as well, but the temporal lobe seems to house at least two functionally distinct rhythms. One of these is the *auditory tau rhythm*, which is generated in the posterior/middle section of the superior temporal gyrus [68, 41]. It is still controversial if the auditory tau rhythm can be observed in resting-state MEG recordings [42]. Besides the tau rhythm, the temporal lobe generates at least one other alpha rhythm, known as the *breach rhythm* or the *third rhythm* [11, 57]. It is located over the midsection of the middle/inferior temporal gyrus and in contrast to the tau rhythm, is not modulated by auditory stimuli [68, 42, 70]. Due to its relatively low frequency (6–9 Hz) some studies have classified it as theta (4–8 Hz) [22, 31]. It is unclear how the breach rhythm relates to the auditory tau rhythm and if the two are actually distinct.

The contours of the organization of spontaneous alpha in the human cortex has been obtained by piecing together observations of different data-sets and recording techniques (MEG/EEG/ECoG). Outstanding questions are if the known alpha rhythms are present in the majority of people, how large their intra- and inter-subject variability is, how many more generators exist in the resting human cortex, and how the generators are coordinated to form resting-state networks (RSNs). We address these questions using the unique MEG data-set provided by the Human Connectome Project (HCP) (MEG2 Data Release) [39, 69]. This data-set contains three resting-state sessions from a large number of (healthy young) participants (N = 94) and have employed state-of-the-art surface-based cortical registration techniques developed by the HCP [20].

In Section 2.1 we analyze the cortical distribution of power, spectral, and temporal features of alpha and conduct a test-retest reliability analysis. We find that power maps are reliable to the extent that they can be used to identify individual subjects from the population. Frequency maps, on the other hand, are found to be more sensitive to momentary (i.e. within-scan) cortical state and might therefore explain some of the inter-scan variability in cognitive, perceptual, and emotional variables. In Section 2.2.1 we describe how non-negative matrix factorization (NMF) can be used to delineate cortical generators. In Section 2.2.2 we apply NMF to spontaneous power fluctuations and identify 16 reliable generators in each hemisphere, which include the classical alpha rhythms in visual, somatosensory, and auditory cortices, as well as additional ones in the ventral occipital and medial temporal lobes, inferior parietal lobule, and the parieto-temporal junction. We also found a second generator within the primary somatosensory cortex, located more ventrally w.r.t. the classical mu rhythm. No generators were found in the frontal cortex. In Section 2.2.3 we show that in addition to the generator locations, their relative strengths are also reliable, that is, reproducible across scans. In Section 2.2.4 we show that the generators’ power fluctuations are coordinated across hemispheres and hence form resting-state networks (RSNs). Our study thus demonstrates that cortical alpha is organized into a multitude of homologue RSNs and, as such, provides the most comprehensive characterization to date of the organization of alpha in the resting human cortex.

## 2 Results

### 2.1 General features of resting-state cortical alpha

#### 2.1.1 Power and frequency maps

In this section we investigate the cortical distribution of power and oscillatory frequency. For each subject and run, we averaged the power matrix over epochs, obtaining a single vector (or map) that contains the average power for all cortical vertices (see Materials and Methods). The power maps where subsequently divided by their sum to obtain subject- and run-specific power-density maps. This was done to remove global power differences between subjects that might dominate the analysis. Subject- and run-specific frequency maps were also obtained (see Materials and Methods). Global intra-subject (inter-run) and inter-subject were removed by subtracting the subject- and run-average frequency from each frequency map. This hence yielded subject- and run-specific relative frequency maps.

Figure 1A shows the subject- and run-averaged power-density map. Alpha power is particularly high in the occipital and temporal lobes, and to a lesser extent in parietal lobes, extents into the central sulcus, and is particularly low in the frontal cortex. Roughly the same distribution has been reported in [31], using a different MEG scanner and a different source projection method. Figure 1B shows the subject- and run-averaged relative frequencies. Although the frequency range is rather narrow (±0.35 Hz), the frequency difference between posterior and frontal regions in individual subjects can be much larger (up to 2 Hz). Particularly high frequencies can be observed in occipital and parietal midline regions, where the classical posterior alpha rhythm is located [27, 10, 46] as well as on the post-central gyrus, where the classical somatosensory (mu) rhythm is located [27, 64, 63, 46]. Note also that the frequency is particularly low in (medial) frontal cortex. Because the frequencies were calculated not as peak-frequencies but as the center frequencies using a weighted average over frequency bins within the alpha-band (see Materials and Methods), these low frequencies are due to the absence of alpha power in frontal cortex (see Figure 1A), and therefore predominantly reflect frontal theta activity, which is known to be generated here [32, 65].

**Figure 1:**
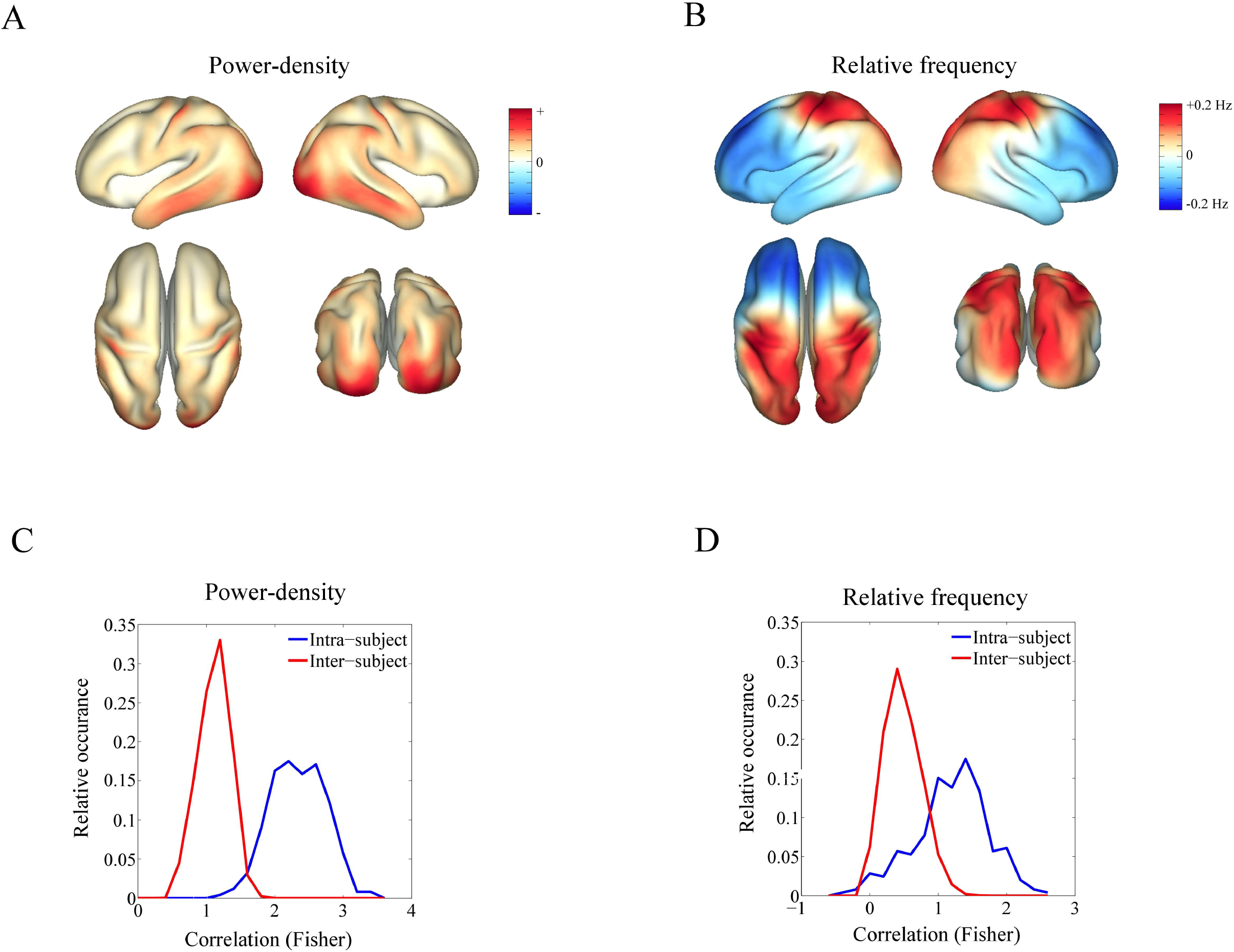
Cortical power and frequency maps. A. Color-coded power-density map, averaged over runs and subjects. The values are non-negative and sum to one. B. Color-coded relative frequency maps, averaged over runs and subjects. The colorbar ranges from −0.2 Hz to 0.2 Hz. C. Intra- and inter-subject distributions (blue and red, resp.) of the Fisher-transformed spatial correlations between the relative power maps. D. Intra- and inter-subject distributions (blue and red, resp.) of the Fisher-transformed spatial correlations between the frequency power maps.

#### 2.1.2 Test-retest reliability

To compare intra- and inter-subject variability of the power and frequency maps, we calculated the distributions of the intra- and inter-subject spatial correlations. The intra-subject distribution was obtained by calculating, for each subject, the spatial correlation between the power maps of the three unordered pairs of runs and pooling the correlations from all subjects. Spatial correlations were calculated as the cosine of the angle between the respective maps. The inter-subject distribution was obtained by calculating, for each run, the spatial correlation between all ordered pairs of subjects and pooling to correlations from all runs. The resulting intra- and inter-subject distributions of the (Fisher transformed) power-density correlations are shown in Figure 1C. The fact that almost all intra-subject correlations are higher than all inter-subject correlations implies that these maps can be used as functional fingerprints in that they allow a given subject to be identified from the group of all 82 subjects if its power-density map on one of the runs is given. This is in analogy with resting-state BOLD-fMRI functional connectivity matrices, which have recently shown to posses this property [18]. In contrast, the frequency maps cannot be used as functional fingerprints mostly because of their relatively high intra-subject variability (in the sense of low spatial correlations) as can be seen from Figure 1D.

Figure 2A shows the power-density maps of two subjects (with HCP identification numbers 112920 and 990366) for each of the three runs and displayed on their native cortical meshes. For both subjects, the power-density maps are rather similar across runs (low intra-subject variability) and are relatively different between the two subjects (high inter-subject variability). It is the combination of these two properties that enables identification of individual subjects. In this example, the first subject (ID 112920) has a relatively strong posterior alpha rhythm while the second subject (ID 990366) has a relatively strong somatosensory rhythm in the left hemisphere and a strong rhythm in the supramarginal gyrus in the right hemisphere. Figure 2B shows the relative frequency maps for the same two subjects. These maps are clearly more variable between runs as compared to the power maps. In particular, the relative frequency of the posterior alpha rhythm in the first subject is higher in run 2 as compared to run 1 and still higher in run 3. An interesting observation is that while the power maps of the first subject do not clearly show the presence of the somatosensory rhythm, the frequency maps show a locally increased relative frequency around the post-central sulcus, particularly in the first two runs. The mu rhythm of the first subject is hence definitively present, but very weak, which makes it hard to discern it in the power map. In Section 2.2 we show that such weak generators can still be detected by an appropriate decomposition of the power matrices.

**Figure 2:**
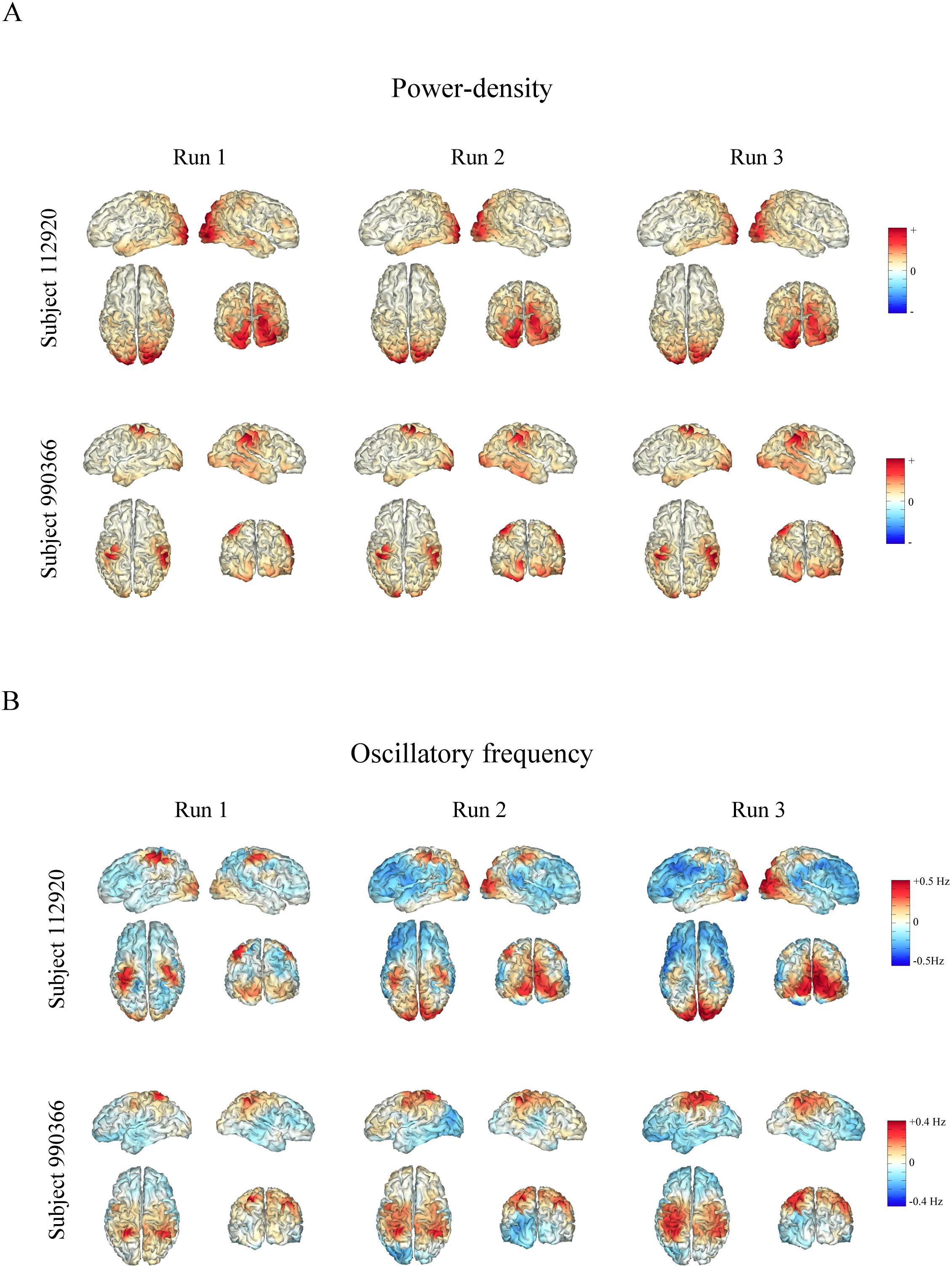
Power and frequency maps in two subjects. A. Power-density maps for two HCP subjects (ID 112920 and ID 990366) for each of the three resting-state runs. The colors are scaled independently for each subject and run (maximal range). B. Relative frequency maps for the same subjects. For subjects 112920 and 990366, the colorbar ranges between ±0.5 Hz and ±0.4 Hz, respectively.

#### 2.1.3 Temporal dynamics

To characterize the temporal dynamics of spontaneous low-frequency (< 0.5 Hz) power fluctuations within the alpha-frequency band, we calculated, for each cortical vertex and run, the subject-averaged spectral densities. To have a shared frequency resolution across subjects, we used the lowest number of epochs in the entire data-set (110), which corresponds to a fundamental frequency of 0.0045 Hz. The highest observable (Nyquist) frequency is determined by the sampling frequency (0.5 Hz) and equals 0.25 Hz. To characterize the power densities, we calculated their Hurst exponents. For a spectral density of the form 1/*f^α^*, the Hurst exponent α measures the slope of the log-transformed spectral density and hence characterizes the power-law scaling of the amplitude fluctuations [43]. Vertex-wise Hurst exponents were calculated by fitting first-order polynomials to the log-transformed spectral densities. An overall Hurst exponent was also calculated by fitting a polynomial to the vertex- and run-averaged power spectral densities. This yielded a Hurst exponent of 0.32. Figure 3A shows the overall power spectral density, together with the fitted log-linear polynomial. It shows that the power spectral density is well-modeled by a power-law and hence is scale-free, at least over the range of observable frequencies (two orders of magnitude). The Hurst exponents obtained from the run-specific spectral densities (so before averaging over runs) were equal to 0.30, 0.32, and 0.35, for run 1, 2, and 3, respectively. Moreover, the average spatial correlation of the Hurst maps over the three ordered run-pairs equals 0.83. These observations shows that the Hurst exponent is a robust and reproducible feature of resting-state cortical dynamics within the alpha frequency-band, at least on the time-scale of the experiments (several hours). Figure 3B shows the overall cortical Hurst map. Note the relatively low exponent in (medial) frontal cortex. Thus, besides power and frequency, frontal alpha-band activity seems to behave differently from other cortical regions also with respect to its temporal dynamics.

**Figure 3:**
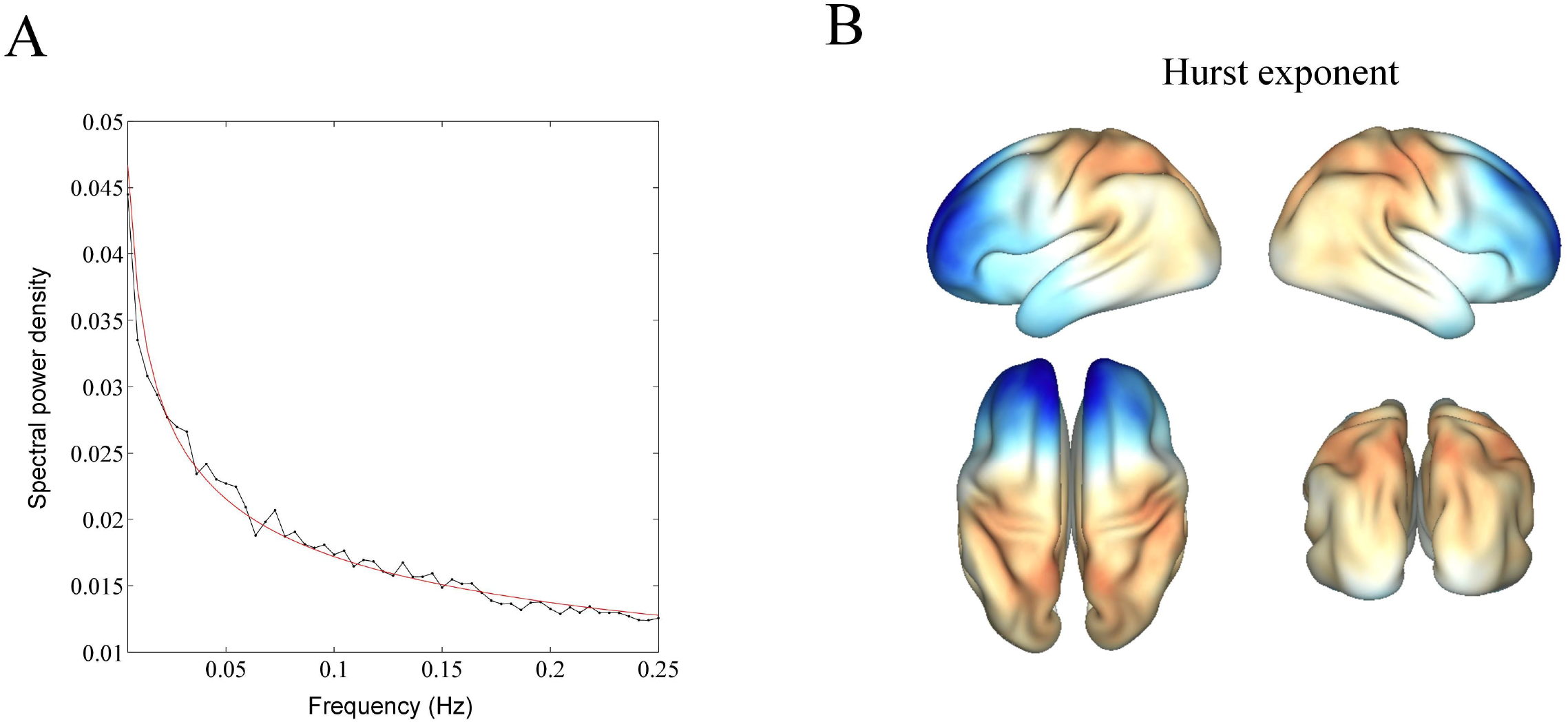
Temporal dynamics of low-frequency power fluctuations. A. Overall power spectral density of epoch-wise source power, obtained by averaging the vertex- and run-wise spectral power densities (black line). The red line corresponds to the fitted log-linear polynomial and has a Hurst exponent of 32. B. Cortical Hurst map obtained by averaging the subject-averaged Hurst exponents over runs and cortical vertices. The exponents range from 0.15 to 0.40.

### 2.2 Generators of resting-state cortical alpha

In the next section, we use non-negative matrix factorization (NMF) [6] of the low-frequency (< 0.5 Hz) power fluctuations to decompose them into a number of generators (Section 2.2.1). NMF decomposes the power matrices into a small number of non-negative spatial and temporal components that together account for the spatiotemporal structure present in the fluctuations, and, as such, uncovers latent structure in their spatiotemporal dynamics. We subsequently compare the generators with known generators from classical MEG dipole studies (Section 2.2.2), consider their contribution to cortical power (Section 2.2.3), and characterize their organization into resting-state networks (Section 2.2.4).

#### 2.2.1 Delineation using non-negative matrix factorization

To break down the spontaneous low-frequency (< 0.5 Hz) alpha-band power fluctuations into generators, we used non-negative matrix factorization, also known as parallel factor analysis (PARAFAC) [6]. For each subject *s* (*s* = 1, ⋯, 82) and run *r* (*r* =1, 2, 3), the *n* × *m_s,r_*-dimensional power matrix *P_s,r_* was calculated, where *n* denotes the number of cortical vertices (*n* = 8004) and *m_s,r_* denotes the number of epochs of subject *s* and run *r* (see Materials and Methods for the calculation of the power matrices). Subsequently, the *n* × *m_r_*-dimensional matrix *P_r_* is constructed by horizontally concatenating the matrices *P*_1,*r*_, ⋯, *P*_82,*r*_, where *m_r_* denotes the total number of epochs on run *r*. To be sensitive to weak generators, we factorized the power-density matrix 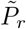, which is obtained from *P_r_* by normalizing its columns by the sum of their entries. A factorization of 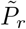, however, induces a factorization of *P_r_*, with the same spatial components, but with scaled time-courses (see Materials and Methods). The power-density matrix 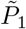of run 1 is shown in Figure 4.

**Figure 4:**
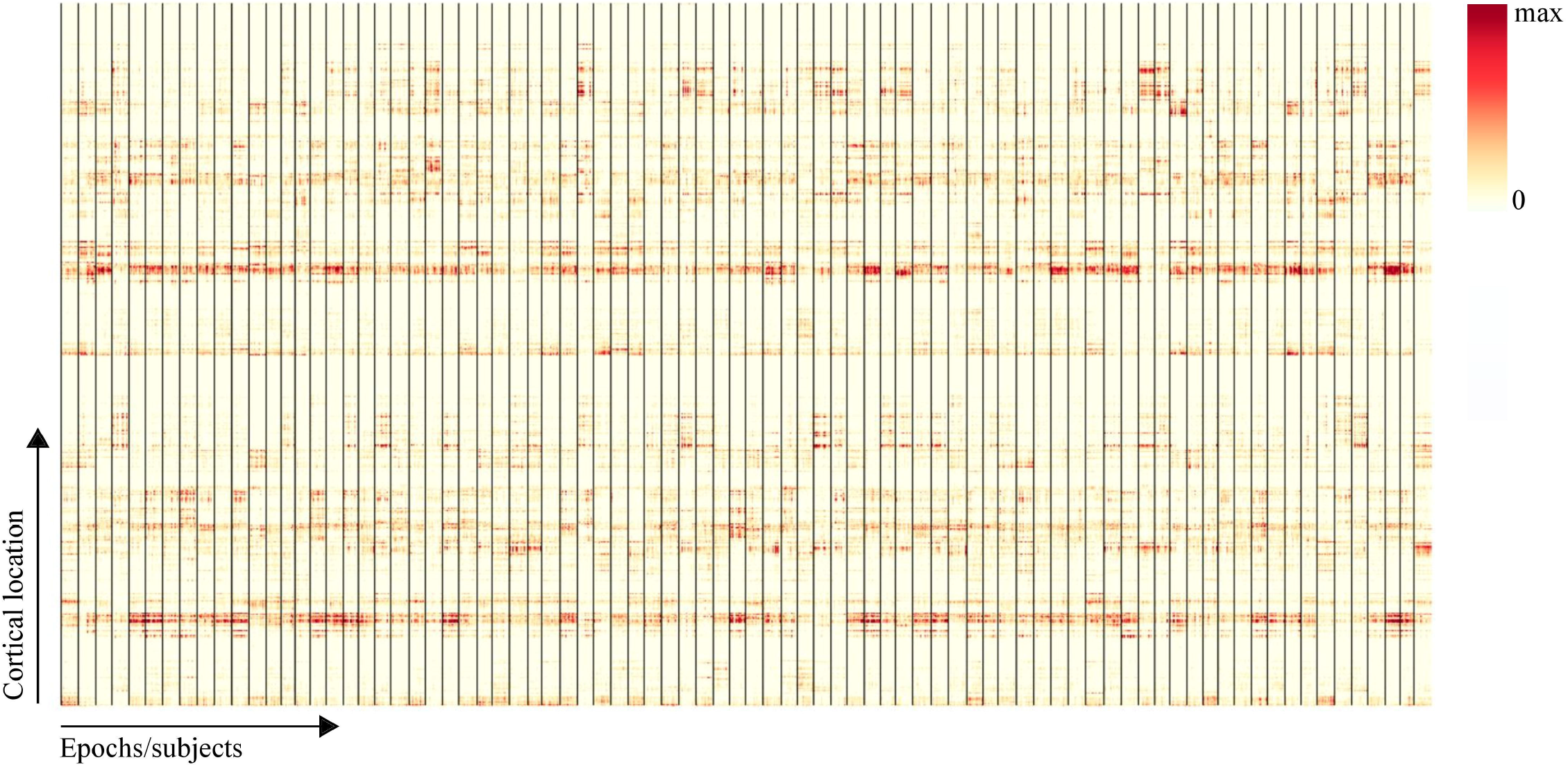
Power-density matrix. Shown is the *n* × *m*_1_-dimensional power-density matrix 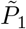 of run 1, which is obtained by concatenating the subject-specific power-density matrices 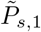 for *s* = 1, ⋯, 82 of run 1. *n* denotes the number of cortical vertices (*n* = 8004) and m\ denotes the total number of epochs on run 1. Subjects are separated by vertical black lines.

We applied the factorization separately to the left and right hemispheres. This was done because when factoring both hemispheres simultaneously, the algorithm yields both lateralized and homogolue components, which complicates subsequent analysis. Our strategy is to first break down the power fluctuations into elementary generators, and subsequently assess their organization into functional networks. Factorization of the left- and right-hemispheric power matrices 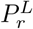 and 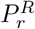 of run *r* yields approximations

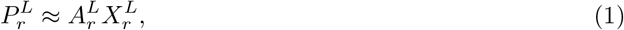

and

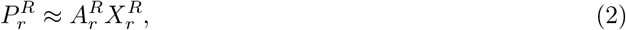

where 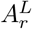 and 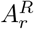 are *n*/2 × *k*-dimensional non-negative matrices that contain the left- and right-hemispheric spatial components (generators) in their columns, 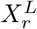 and 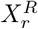 are *k* × *m_r_*-dimensional non-negative matrices that contain the respective time-courses in their rows, and *k* denotes the number of components. To break down the spontaneous power fluctuations into elementary generators, the number of components *k* was set to a large value. Principal component analysis (PCA) of the left- and right-hemispheric power-density matrices showed that *k* = 18 will be sufficient. Crucially, we factorized *P_r_* separately for *r* = 1, 2, 3 and selected only the reliable generators for further analysis, where reliability was defined as a spatial correlation > 0.8. This yielded 17 generators in each hemisphere (and for each of the three runs).

One of the generators had a rather diffuse spatial profile and was confined to the frontal cortex. Given the relatively low oscillation frequencies in frontal cortex (Figure 1B), this generator likely captures the high-frequency end of the frontal theta rhythm and we will not further consider it in this study. The remaining 16 left- and right-hemispheric generators were paired by using the inter-hemispheric spatial correlation matrix of the run-averaged components: Each left-hemispheric generator was paired with that right-hemispheric generator with which it was maximally correlated. The (reliable and paired) run-averaged generators are shown in figure 5.

**Figure 5:**
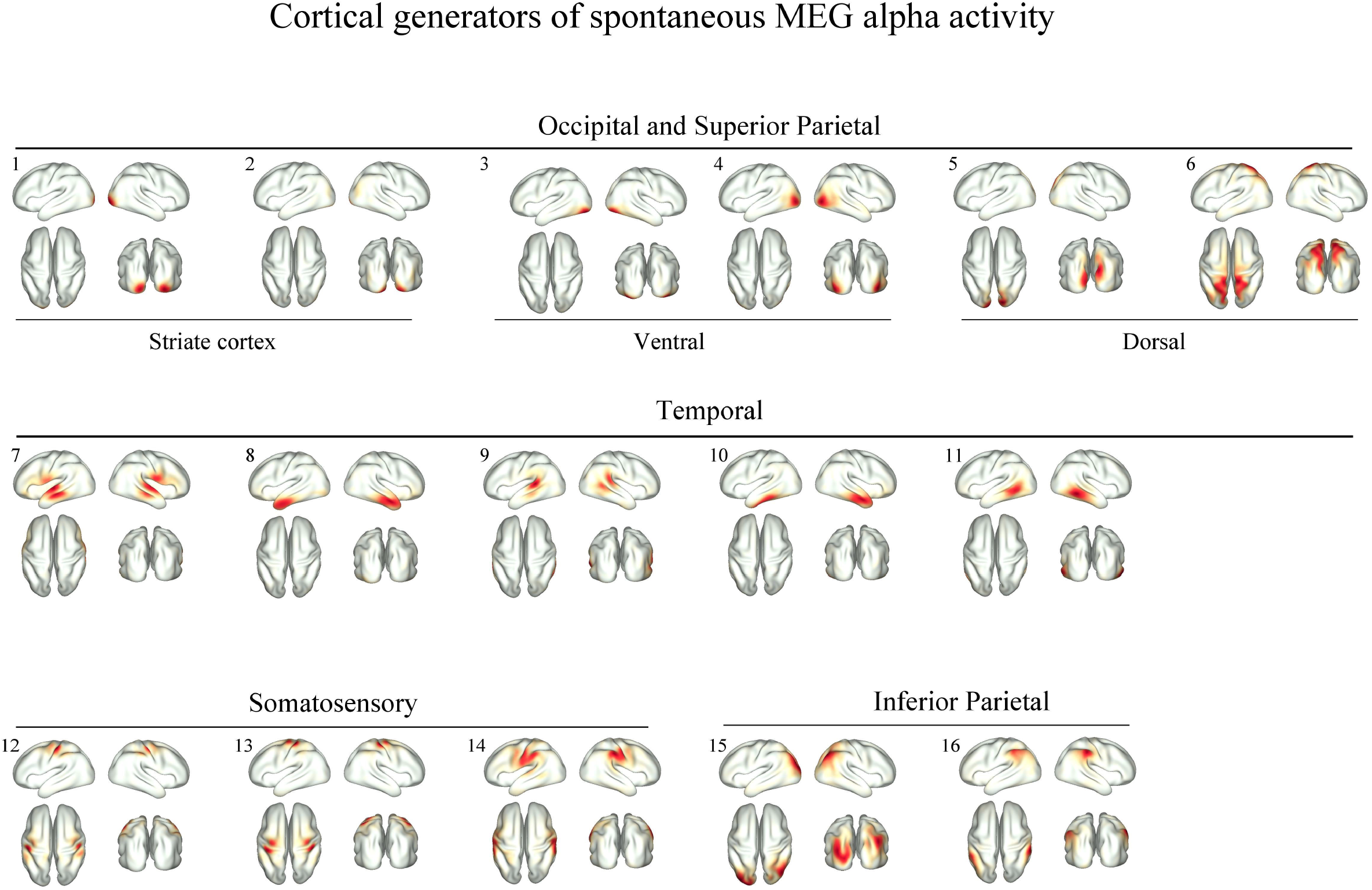
Cortical generators of resting-state MEG power fluctuations within the alpha frequency band. Shown are the run-averaged generators that were reliable, that is, present in all three runs (spatial correlation > 0.8). In each hemisphere, 16 reliable generators were identified, and in this figure they have been grouped based on maximal inter-hemispheric spatial correlation. The generators are ordered according to anatomical/functional location. Colormaps have been scaled to their maximal range.

#### 2.2.2 Relation to the classical alpha rhythms

In this section we link the identified generators to the cortical sources of alpha oscillations as reported in classical MEG dipole studies. Six of the 16 generators are located within the visual cortex. They are numbered 1 to 6 in Figure 5. Generators 1 and 2 are located in striate cortex, around the tip of the Calcarine sulcus, generators 3 and 4 are located in ventral visual cortex, and generators 5 and 6 are located in dorsal visual cortex, with generator 6 extending into medial parietal regions. The striate and dorsal generators cover the area where the classical posterior alpha rhythm has been localized in early MEG studies using equivalent current dipoles (ECDs), that is, around the tip of the Calcarine sulcus and around the parieto-occipital fissure [27, 10, 46]. In addition to these classical generators of the posterior alpha rhythm, the ventral generators have less frequently been reported [45]. The factorization thus has broken down the classical posterior alpha rhythm into several localized generators with partially independent time-courses. This is reminiscent of the results of these classic studies, which reported a multitude of independent ECDs, scattered throughout the Calcarine sulcus and the parieto-occipital fissure [27, 10, 46]. Our observations hence corroborate these findings and reinforce the notion that the posterior alpha rhythm is not a unitary rhythm, but a spatially extended system of oscillations [72].

Three of the 16 generators are located within the somatosensory cortex. They are numbered 12 to 14 in Figure 5. Generator 13 is located on the post-central gyrus and corresponds to the *somatosensory mu rhythm*, which originates from the hand representation area of SI [64, 63, 46]. The weaker coverage of the pre-central gyrus is due to the inability of the source projection method to accurately reconstruct sources located of the fundi and walls of sulci (surface bias). The true location of generator 13 hence most likely is on the post-central wall. Generator 12 is also located on the post-central gyrus, but more laterally. This generator might correspond as well to the mu rhythm, reflecting inter-subject differences in the location and spatial extent of the hand representation area. The existence of more than one mu generator suggests that like visual alpha, the mu rhythm is a spatially extended and heterogeneous phenomena, in line with earlier MEG dipole studies [46]. We will refer to generator 13 and 12 as *dorsal* and *ventral mu*, respectively. Generator 14 seems to originate from the secondary somatosensory cortex (SII) and hence corresponds to the *sigma rhythm* [64, 56, 42]. We suspect that this is the same generator as observed in [42], which was localized close to the mouth area of SI.

Five out of the 16 generators are located within the temporal lobe. They are numbered 7 to 11 in Figure 5. Generator 7 covers the middle section of the superior temporal sulcus and generator 8 and 10 cover the anterior section of the middle and inferior temporal gyrus (but not the temporal pole). Given their locations, and the relatively low frequency of alpha in these regions (see Figure 1B), we suspect these generators to correspond to the classical *breach rhythm* [11] also called the *third rhythm* [57], which has been recorded with epidural electrodes [11, 57]. Generator 9 covers the posterior section of the superior temporal sulcus and can hence be identified with the classical *auditory tau rhythm* [68, 41]. Generator 11 covers the posterior section of the middle temporal gyrus and as far as we know, there are no reports on spontaneous MEG/EEG alpha-band oscillations originating from this region.

Two of the 16 generators are located within the inferior parietal cortex. They are numbered 15 and 16 in Figure 5. Generator 15 covers the supramarginal gyrus. Observe, however, that in the left hemisphere, generator 15 also covers medials parts of the occipital lobe. We suspect that this generator can be broken down further into more elementary generators. Generator 16 covers the angular gyrus. We are not aware of any reports of alpha generators in these regions from resting-state MEG (or EEG) data.

#### 2.2.3 Relative contributions

In this section we consider the generators’ relative contributions to the total cortical power. We illustrate this by using the ventral mu rhythm (see Figure 5) but similar observations can be made for the other generators. For each of the 16 generators, we calculated their subject- and run-specific relative contributions 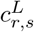(i) and 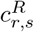(i) to the power-density, where *i* indexes the generators. Thus, 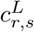(*i*) and 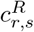(*i*) range between zero and one and their sum over *i* equals one. As an example, Figure 6A shows the population profiles of the relative contributions of the left-hemispheric ventral mu rhythm (generator 12) for all subjects and all three runs. The relative contributions of the ventral mu rhythm vary considerably between subjects, but are reproducible over scans. The same holds for the other generators. We quantified the reproducibility of the relative contribution of a given generator by calculating, for each of the three ordered run-pairs, the average correlation between the population profiles and compared that with the respective correlation values obtained by first randomly permuting the subject numbers. For all generators and both hemispheres, none of the 10^4^ null-values exceeded the respective observed values, which yielded a *p*-value of *p* < 0.0032, after correcting for multiple (2 × 16 = 32) comparisons. This shows that the high test-retest reliability of the cortical power maps (see Section 2.1.2) is inherited from the high reliability (together with their high variability over subjects) of the relative contributions of the underlying generators.

**Figure 6:**
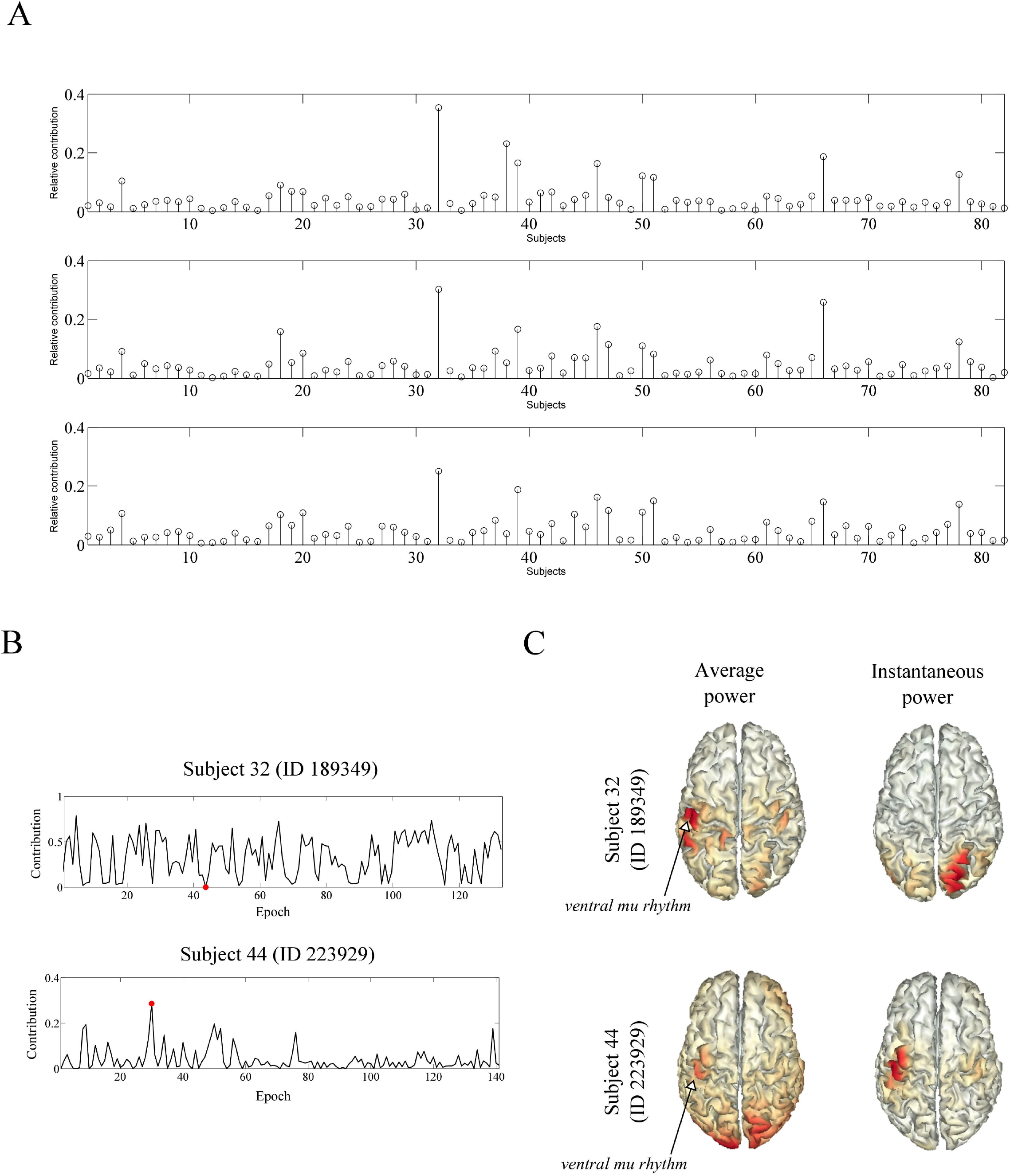
Relative contributions of the ventral mu rhythm. A. Relative contributions of the left-hemispheric ventral mu rhythm (generator 12) for all subjects and all three runs. B. Instantaneous (epoch-wise) relative contributions of the ventral mu rhythm for two subjects Top and bottom panels correspond to subject 32 (ID 189349) and subject 44 (ID 223929), respectively. C. Power maps (left column) and instantaneous power maps (right columns) of the same two subjects. The instantaneous power maps of subject 32 and 44 were taken at epoch numbers 44 and 30, respectively, which are designated by the red dots in B.

The indices 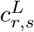(i) and 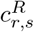(*i*) measure the relative contributions of the *i*-th generator over the entire run, and as such, do not provide information on their within-run temporal fluctuations. Information on the latter can be obtained by calculating the instantaneous (that is, epoch-wise) relative contributions (see Materials and Methods). Figure 6 shows the instantaneous relative contribution of the ventral mu rhythm in two subjects. Subject 32 (ID 189349) and subject 44 (ID 223929) have a relatively large and small average relative contribution of the power-density (see Figure 6A). This is reflected in their (average) power maps, which are shown in Figure 6C (left column). As is clear from Figure 6B, in both subjects, the instantaneous relative contribution of the ventral mu rhythm is far from being constant. While in subject 32 the ventral mu rhythm is dominant on average, there are several epochs during which its (instantaneous) contribution is close to zero. The reverse is true for subject 44: Although its average contribution is low, there are epochs during which the ventral mu rhythm is relatively dominant. This is illustrated in Figure 6C (right panel), which shows the instantaneous power maps of these two subjects, taken at epoch numbers 44 and 30, as designated by the red dots in Figure 6B. Similar observations can be made for the other generators.

#### 2.2.4 Organization into functional networks

To assess the inter-hemispheric correlation structure of the identified generators, we correlated the generators’ time-courses with the low-frequency power fluctuations of each cortical vertex in the contra-lateral hemisphere. This yielded an inter-hemispheric correlation map for each of the left- and right-hemispheric generators and for each of the three runs, which were subsequently rectified to retain only the non-negative correlations (see Materials and Methods for details). The run-averaged rectified correlation maps are shown in Figure 7. The run-specific correlation maps were reproducible over runs: their spatial correlation, averaged over each of the three combinations of ordered run-pairs and over generators, was 0.91.

**Figure 7:**
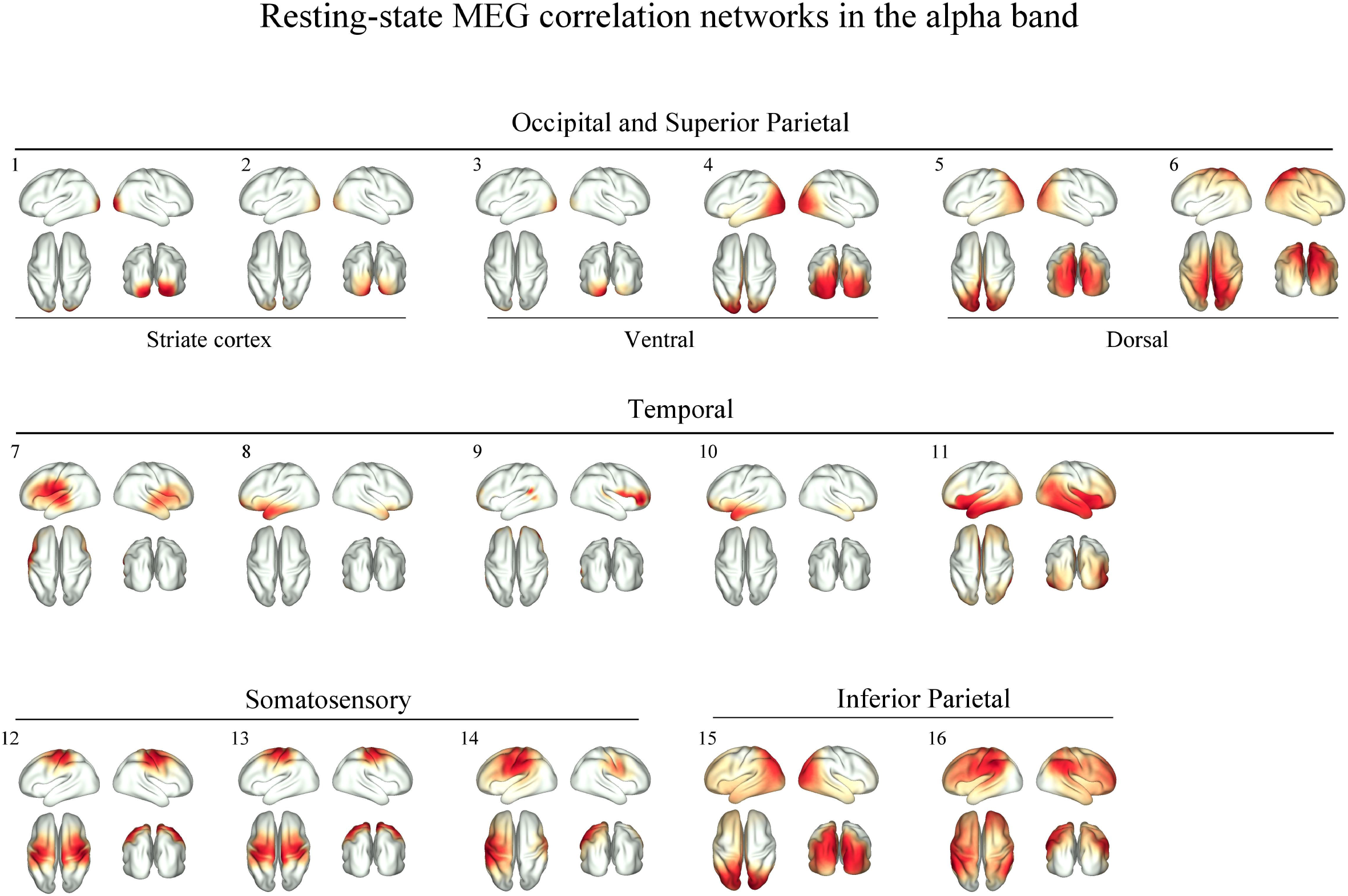
Resting-state functional networks within the alpha frequency band extracted using correlations. Shown are the run-averaged rectified inter-hemispheric correlation maps of the 16 identified generators. Their ordering is identical to that of Figure 5. Colormaps have been scaled to their maximal range.

For each generator, the highest correlations are found roughly at the respective homologue cortical regions and this holds for both the left- and right-hemispheric generators. This means that the generators’ spontaneous power fluctuations are coordinated across hemispheres and the generators thus are organized into homologue functional networks. Note that for most generators, the correlation maps are less localized than the corresponding spatial components (Figure 5), which shows that functional correlations are relatively sensitive to magnetic field spread. The lack of localization is particularly severe in cortical regions that have weak MEG leadfields like the insular cortex, the temporal pole, and medial regions. This sensitivity to magnetic field spread can be alleviated by using covariances instead of correlations [25]. This is shown in Figure 8. The reason why the covariance structure is more localized than the correlation structure is that covariances incorporate the amplitudes of the time-courses and amplitudes are less susceptible to magnetic field spread in the case of MEG or volume-conduction in the case of EEG [29]. A potential drawback of using covariances over correlations, however, is that functional coupling of a generator to a low-amplitude region can be overlooked. This is the case for the ventral attention network (VAN) and the default mode network (DMN) as we will see below.

**Figure 8:**
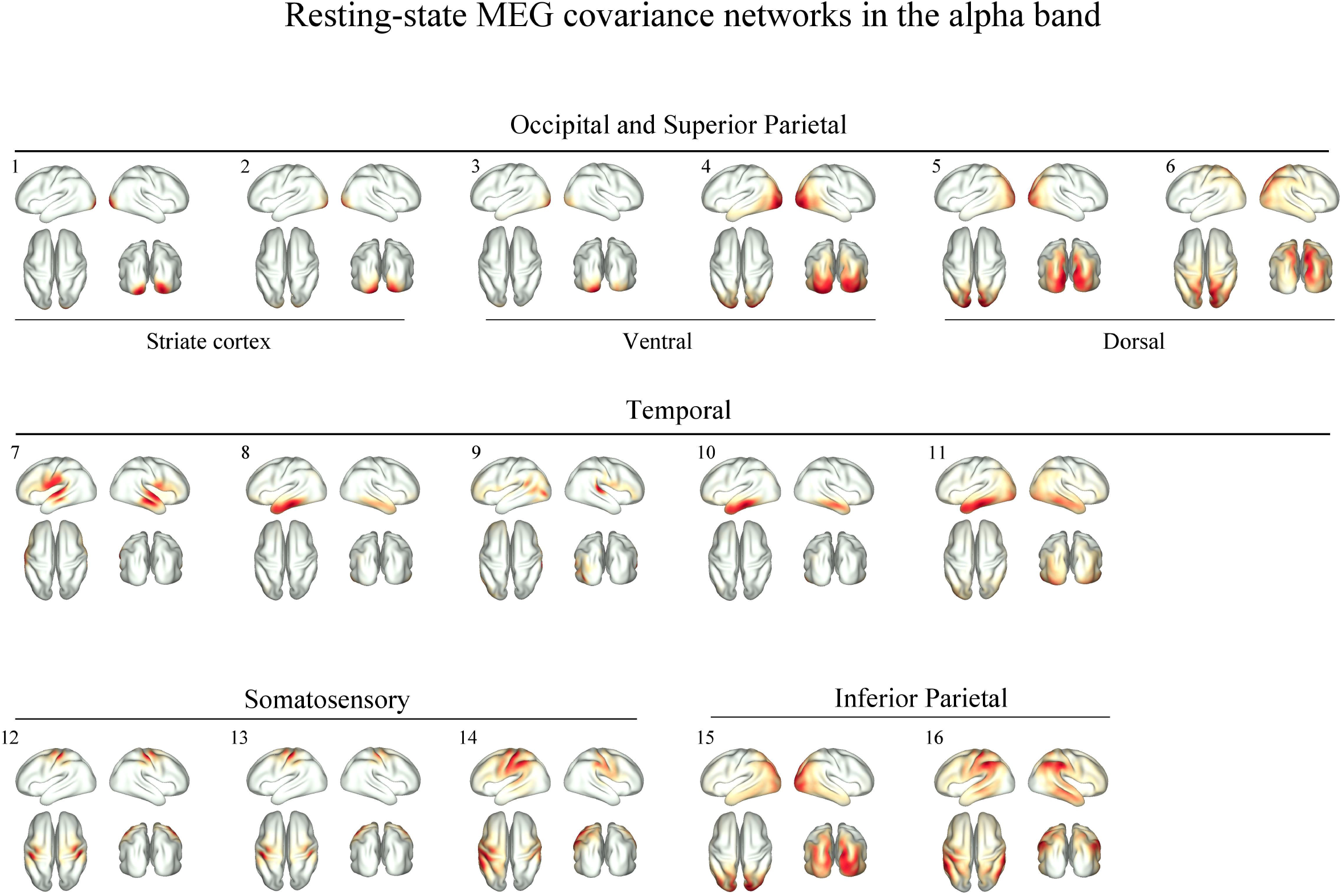
Resting-state functional networks within the alpha frequency band extracted using covariances. Shown are the run-averaged rectified inter-hemispheric covariance maps of the 16 identified generators. Their ordering is identical to that of Figure 5. Colormaps have been scaled to their maximal range.

The networks of most generators are simple, in the sense of comprising a single region in the contralateral hemisphere. Exceptions, however, are the networks formed by generators 9 and 16 (Figure 7). Generator 9, which we earlier identified with the tau rhythm of the temporal lobe, is functionally connected to inferior frontal cortex, which is particularly clear for the right hemisphere. We suspect this generator to be part of the VAN, which comprises inferior frontal cortex and the temporal-parietal junction, is known to be right-lateralized [14], and is part of the task-positive network, which is known to operate within the alpha-frequency band [37]. Because frontal regions have relatively low alpha-power (Figure 1A), functional connectivity with frontal regions is suppressed when using covariances instead of correlations, although still observable (Figure 8). The same holds for generator 16, which is located on the supramarginal gyrus and is functionally connected to medial prefrontal cortex (Figure 7). These regions form, respectively, the parietal and frontal nodes of the DMN, which has been observed before within the alpha-frequency band [37]. Frontal connectivity is suppressed when using covariances (Figure 8), although the connectivity to the temporal node of the DMN is better visible due to relatively strong alpha oscillations within the temporal lobe (Figure 1A). Generally, therefore, the most complete picture of the functional organization of cortical generator is obtained by using both correlation and covariance maps.

## 3 Discussion

### Main findings

In this study we have used the resting-state MEG data-set provided by the Human Connectome Project (HCP) [39], which comprises a large number of subjects (*N* = 94) and three scans per subject, to assess the functional organization of human resting-state cortical activity within the alpha frequency band (7–14 Hz). We have used a non-negative spatiotemporal factorization [6] of source-projected power fluctuations to identify reliable population-level alpha generators. We found 16 generators in each hemisphere which include the well-known alpha rhythms in visual, somatosensory, and auditory cortices [68, 64, 41, 56, 46, 42], as well as additional generators in temporal and inferior parietal cortices, which, to the best of our knowledge, have not been reported in previous studies. No generators were found in the frontal cortex. By correlating their power fluctuations with those at all cortical locations, we found that the generators are coordinated across hemispheres, thus forming resting-state networks (RSNs). Our study hence uncovered a multitude of RSNs within the alpha band, including visual, somatosensory, and auditory subnetworks, the default mode network (DMN) and the ventral attention network (VAN) and demonstrated that these networks are supported by the classical alpha rhythms. These findings provide a more complete and detailed picture of the organization of resting-state cortical alpha and illustrate the power of the HCP MEG data-set.

### General features of cortical alpha

Cortical alpha power was strong throughout most of the post-central cortex and particularly in visual, somatosensory, and temporal regions, where the classical alpha rhythms reside [27, 10, 46]. Frontal (i.e. pre-central) regions had relatively low power. Strikingly similar power maps have been reported before, using a different MEG scanner and a processing pipeline [31], which demonstrates their robustness. Our findings and those of [31] indicate that the frontal cortex doesn’t generate alpha. Although frontal alpha has frequently been reported in EEG and MEG studies, peak-frequencies are typically lower than those of more posterior regions [1, 31]. This is clearly illustrated in Figure 1B. Combined with the fact that we found only one frontal generator - which was diffuse and therefore likely reflects a deep generator - frontal alpha might perhaps be better classified as theta, which is known to be generated within the (medial) frontal cortex and whose frequency range overlaps with that of alpha [32]. An additional argument for the conflation of frontal alpha and theta is that we extracted alpha oscillations based on subject-specific alpha peak-frequencies: Had we used a standard frequency band, which is common in EEG and MEG studies, perhaps we would have identified more frontal generators (and classified them as alpha). We therefore suspect that the frontal generator observed in this study, reflects the high-frequency end of the frontal theta rhythm. A similar consideration might apply to the anterior part of the temporal lobe [57, 68, 41] and refining our taxonomy of resting-state cortical alpha/theta will be an important step in further understanding their functional organization. Concerning temporal features of cortical alpha, we found the source-projected power fluctuations within the alpha frequency band obey power-law scaling over two-orders of magnitude (from 0.01 to 0.25 Hz) with Hurst exponents displaying a posterior-anterior gradient. These findings essentially reproduce earlier ones in sensor-level MEG data [43, 53] and indicate that local cortical circuits involved in the generation of alpha operate in the vicinity of a second-order phase transition [43].

### Test-retest reliability

Cortical power maps were highly reproducible across scans and differed substantially between subjects. Such a combination of low intra-subject (i.e. inter-scan) and high inter-subject variability allows these maps to be used as functional fingerprints to identify single subjects from a population. This finding is not new, as several studies have demonstrated that scalp EEG power maps are reliable over even longer periods [38, 17]. In contrast, the test-retest reliability - which is quantified by the ratio of intra-subject and inter-subject variability - of cortical frequency maps was lower and this could be almost entirely contributed to a higher intra-subject variability. Although this implies that frequency maps are less useful for subject identification, albeit only marginally as shown before for EEG [55], it is precisely the high intra-subject variability that enables their use as a marker for momentary (i.e. within-scan) cortical state. Inter-trial variability of a variable usually reflects statistical noise and is therefore of no interest. In the case of oscillatory frequency estimated from five minutes of data, statistical noise is practically absent, and hence the intra-subject variability in frequency maps reflects actual variability. This makes frequency maps sensitive to momentary cortical state and might explain some of the intra-subject variability in behavioral, cognitive, perceptual, or emotional state [16]. Note that intra-scan variability in psychological variables cannot be explained by cortical power maps, as they are practically identical across scans. Low test-retest reliability, together with the absence of statistical noise, thus opens up an interesting way to study intra-subject variability in mental state and again illustrates the (potential) power of the HCP MEG data-set.

### Cortical alpha generators

Early MEG dipole studies have shown that the visual, somatosensory, and auditory cortices generate alpha oscillations in the resting-state [27, 10, 46]. Although there has been some MEG evidence for the existence of additional generators in higher-order sensory cortices [56, 42], they have not been observed in all studies and it remains controversial if they are observable with MEG. In stark contrast, a multitude of generators has been inferred from MEG data by using task-related designs and trial-averaging. These and related studies have established the existence (and functional relevance) of multiple generators in early and higher-order sensory cortices as well as in associative cortex [74, 66, 24, 70, 50, 67]. By only using resting-state MEG data, our findings can be thought of as providing more direct evidence for their existence. They also demonstrate that most (know) alpha generators are observable with MEG without using experimental contrasts or trial-averaging, something which is considerably more challenging from a signal-processing point of view. To delineate the generators, we used a combination of source projection and matrix factorization. Although inverse modeling yields cortical power maps, which show that alpha is widespread throughout the post-central cortex (see Figure 1A and [31]), by themselves, such maps do not allow to delineate different generators. Likewise, although factorization of sensor-level MEG data allows for the identification of generators, without source projection, their locations cannot be inferred. Our results hence illustrate the strength of combining source projection and matrix factorization in elucidating the functional organization of resting-state cortical activity [47, 44].

### Relation to resting-state networks

Resting-state networks in early visual, somatosensory, and auditory cortices in the alpha frequency band have been observed before with MEG [30]. In [30] and several other studies ([7, 13], for example), networks were identified by seed-based correlation maps, which are relatively sensitive to magnetic field spread. As a consequence, the localization of these networks is relatively rough. For instance, placing a seed in the (early) somatosensory cortex, yields correlation maps that cover both the pre- and post-central gyrus over most of their lateral extent. Although magnetic field spread can be mitigated to some extent by using covariances instead of correlations (see Figure 8 and [25]), the factorization approach taken in our study seems to yield even higher accuracy, although the high-quality surface-based cortical registration employed by the HCP might also play a part in this [20]. Another advantage of the factorization approach is that it allows for decomposition of the sensory networks into subnetworks, thereby revealing some of their internal structure. In the visual network, for instance, it allows delineation of early, dorsal, and ventral subnetworks and the same holds for the other sensory modalities. A similar approach identified visual subnetworks from resting-state MEG data, although with lower spatial resolution [3]. By delineating generators prior to a correlation analysis, we have demonstrated that the sensory RSNs in the alpha frequency band are supported by (classical) alpha rhythms and the same holds for the DMN and VAN.

### Functional interpretation

There is causal evidence that resting-state alpha power reflects the excitability of cortical tissue, at least within the visual system [61]. By now, it is established that alpha power is regulated by selective attention and that this has perceptual consequences. This has been shown for the visual [74, 35, 60, 66], somatosensory [15, 24], and auditory [4, 70, 54, 71, 28, 73] modalities as well as across modalities [19, 67]. Viewed in this context, the multitude of alpha generators observed in our study are likely to reflect ongoing fluctuations in the allocation of sensory attention [34, 33]. In addition to sensory generators, we found two generators within the inferior parietal cortex (angular and supramarginal gyrus). Alpha power in these regions has been shown to be modulated during auditory working memory [50] and audiospatial attentional tasks [2]. We observed that the relative strength of the different generators fluctuates during the scanning session. For example, at one particular moment during the scanning session, alpha within the left somatosensory cortex might dominate, while at a subsequent moment, alpha in the right primary cortex dominates. During the resting-state, there thus appears to be a dynamic switching of attentional focus both within and across sensory modalities. In future studies, the delineation of the power fluctuations of individual alpha generator, as enabled by the factorization approach, might proof useful in disentangling separate attentional processes and in characterizing their interactions.

### New applications of the methodology

The methodology proposed in this study, namely, non-negative factorization of source-projected power fluctuations, might be used to address outstanding questions. One of these concerns the electrophysiological signature of cortical generators and the ensuing RNSs. In this study we focused on generators within the alpha frequency band. Previous research has shown, however, that electrophysiological RNSs are generally not confined to a single frequency band [48]. An interesting approach would therefore be to apply such a factorization to the three-way tensor formed by space, time, and frequency triples. This generalization has already been proven useful in identifying separate cognitive processes from scalp EEG recordings [51] and might be equally useful in providing a more general characterization of resting-state cortical generators and RNSs. Another possible application concerns the relation between resting-state EEG alpha powerand BOLD-fMRI. There is currently no general consensus on how BOLD-fMRI relates to cortical alpha [21, 40, 23, 49, 52] and one of the complicating factors is the multitude of cortical alpha generators. In these and related studies, a single generator is usually isolated by selecting a single EEG electrode or an independent component. By applying non-negative matrix factorization to source-projected EEG data, the hemodynamic correlates of separate alpha generators might more easily be disentangled.

## 4 Materials and Methods

### 4.1 Data acquisition

We used the MEG data-set provided by the Human Connectome project (HCP) [69] (MEG2 Data Release) which comprises three six-minute resting-state scanning runs from 94 young healthy participants using the whole-head MAGNUS 3600 (4D Neuroimaging, San Diego, CA) system, housed in a magnetically shielded room. The participants were instructed to remain still (supine position), to relax with eyes open, and to fixate on a projected cross-hair on a dark background. The recorded data were segmented into two-second epochs and pre-processed using a dedicated pipeline that included detection of bad channels and segments, ICA-based artifact rejection (see [39] for details).

### 4.2 Data pre-processing

Prior to source modeling, we discarded the data of those subjects for which alpha-band oscillations could not be detected in one or more of the three runs and subsequently bandpass-filtered the data of the remaining subjects (*N* = 82) about subject-specific alpha-band frequencies. To detect the presence of alpha-band oscillations in the data of subject *s* and run *r*, we first calculated the power-spectra (using the FFT) for each sensor and epoch, averaged the spectra over epochs and sensors, and subsequently searched for a (local) peak in the within the range 6 – 13 Hz. If such a peak was absent in any of the three runs, the subjects’ data were discarded. The sensor time-series of the remaining subjects were subsequently bandpass filtered ±3 Hz about the weighted-frequency 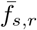

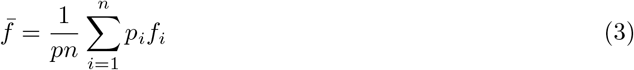

within the alpha-band (6–13 Hz), where *f*_1_, ⋯, *f_n_* denote the frequency bins within the alpha-band, *p*_1_, ⋯, *p_n_* the respective power values, and *p* = *p*_1_ + ⋯ + *p_n_* denotes the total power within the alpha-band. The filtering was done by a fourth-order zero-phase Butterworth bandpass filter. We used 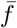 instead of the alpha peak-frequency because the power spectra of many of the subjects contained multiple peaks within the alpha-band. We visually inspected the broadband and bandpass filtered power spectra of all subjects and runs and found the above procedure to well isolate the alpha peaks. After bandpass-filtering, the time-series were down-sampled by a factor of four (from 508.6275 to 127.1569 Hz).

### 4.3 Source modeling

Leadfield matrices were computed in Fieldtrip [58] using realistic single-shell headmodels provided by the HCP. As source spaces we used co-registered individual cortical meshes (4002 vertices per hemisphere) provided by the HCP. A detailed description of the anatomical processing pipeline can be found in [20]. This yielded subject-specific leadfield matrices *G_s,r_* of dimension *m_s,r_* × 3*n*, where *m_s,r_* denotes the number of non-bad MEG channels of subject *s* and run *r* and where *n* = 8004 is the total number of cortical vertices. The columns of the subject- and run-specific leadfield matrix *G_s,r_* thus contain the vertex-wise leadfield vectors for each of the three Euclidean orientations. Suppressing the dependence on subject *s* and run *r* from the notation, the *m* × *t*-dimensional data matrix *B* (where *t* denotes the number of temporal samples) is projected to source space using Harmony [59], which is a smoothed version of the classical minimum-norm estimate (MNE) designed to reduce surface-bias in the source reconstructions. Specifically, the 3*n* × *t*-dimensional source-activity matrix *ĵ* is obtained by applying the Harmony inverse operator 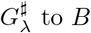 to *B*:

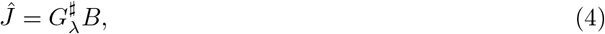

where

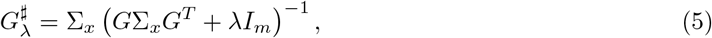

where λ ≥ 0 denotes the regularization parameter, the 3*n*-dimensional matrix Σ_*x*_ the *a priori* source covariance matrix, *I_m_* the *m*-dimensional identity matrix, and where *T* denotes matrix transpose. Harmony models Σ_*x*_ as

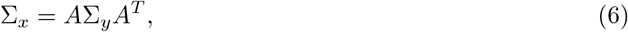

where *A* denotes the 3*n* × 6(*l*_max_ + 1)^2^-dimensional Harmonic transformation matrix, which contains the spherical harmonics (of both hemispheres) up to and including degree *l*_max_ = 20. The 6(*l*_max_ +1)^2^-dimensional matrix Σ_*y*_ denotes the *a priori* source covariance matrix expressed in the Harmonic basis and is modeled as a diagonal matrix with diagonal entries equal to 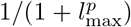, where *p* = 0.5 is a controls the suppression of high-degree Harmonics and therefore the level of spatial smoothing of the source reconstructions [59].

### 4.4 Noise regularization

The regularization parameter λ in Eq. (3) prevents overfitting and choosing an appropriate value is therefore important in obtaining high-quality source reconstructions. For each subject and run, the appropriate level of regularization was determined by applying the *L*-tangent norm criterion [8] to a randomly chosen sample from each epoch and subsequent averaging. The *L*-tangent norm criterion selects that value for λ that minimizes the derivative (i.e. speed) along the (logarithm of the) *L*-curve. Specifically, let

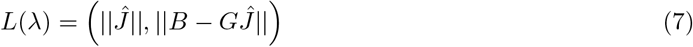

be the *L*-curve [26], then λ is chosen as follows:

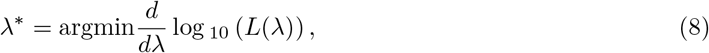

which was obtained by calculating the *L*-curve on the (logarithm of the) interval [−12, −6] in steps of 0.25 and by approximating *d*/*d*λ in these points by finite differences. All obtained values λ* fell well inside the chosen range of λ. Since the samples used for computing the regularization levels were randomly chosen, we repeated their calculation for several subjects to confirm that the same values were obtained. Furthermore, the resulting source-power distributions of all subjects and runs were visually inspected for the presence of over- and under-regularization.

### 4.5 Oscillation frequencies

The a given subject *s*, run *r*, and epoch *e*, and cortical vertex *v*, we calculated the power spectra of the time-series of each cortical vertex (using the DFT), averaged the spectra over all epochs, yielding an epoch-averaged power vector *p* and a frequency vector *f*. The oscillation frequency *f_α_* was then calculated by weighing the frequency vector by the power:

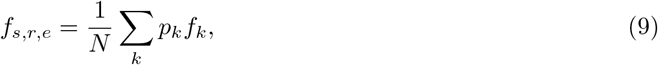

where *N* = Σ_*k*_ *p_k_* is a normalization factor. Note the *f_α_* is not the peak-frequency but corresponds to the center-of-mass of the spectral density *p*.

### 4.6 Power-matrices

For a given cortical vertex, the epoch-wise source-power was obtained from the vector-valued time-course *j*(*t*) = (*j_x_*(*t*), *j_y_*(*t*), *j_z_*(*t*)) by calculating its squared Euclidean norm:

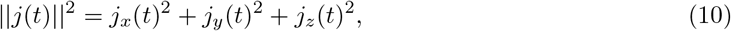

centering its norm to zero by bandpass filtering it ±6 Hz about 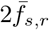, and subsequently averaging the squared samples over the epoch. Centering of ||*j*(*t*)|| removed the (artifactual) low-frequency fluctuations in ||*j*(*t*)|| that resulted from rectifying negative values of *j* by taking its norm. Thus, each subject *s* and run *r* yielded a *n* × *n_s,r_*-dimensional matrix *P_s,r_* of epoch-wise source power, where *n_s,r_* denotes the number of epochs of subject *s* on run *r* and *n* = 8004 is the number of cortical vertices.

### 4.7 Non-negative matrix factorization

Parallel factor analysis (PARAFAC) [6] was applied to the subject-concatenated power-density matrices. To compute the power-density matrices, we first constructed the *n* × *m_r_*-dimensional power matrix *P_r_* by horizontally concatenating the power matrices *P*_1,*r*_, ⋯, *P*_82,*r*_, where *m_r_* denotes the total number of epochs on run *r*:

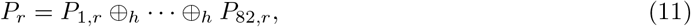

where ⊕_*h*_ denotes horizontal concatenation. Subsequently, the power matrix *P_r_* was converted to a power-density matrix 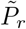. Thus, let Λ_*r*_ be the *m_r_*-dimensional diagonal matrix with *i*-th diagonal entry equal to the sum of the entries of the *i*-th column of *P_r_*, then 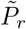 is calculated as

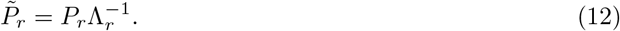

Each row of 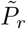 now is a density, that is, its entries sum to 1. PARAFAC is now applied to the left- and right-hemispheric power-density matrices separately. Thus, writing

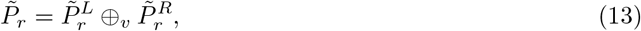

where ⊕_*v*_ denotes vertical concatenation, we factorize 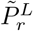 and 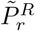 separately, yielding (approximate) factorizations

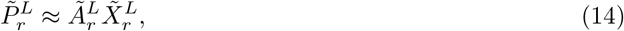

and

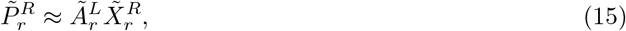

of 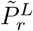 and 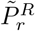 into *k* components, where 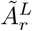 and 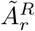 are *n*/2 × *k*-dimensional matrices that contain the spatial components of the left- and right-hemispheric factorizations, respectively, in their columns and 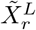 and 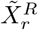 are *k* × *m_r_*-dimensional matrices that contain the temporal components of the left- and right-hemispheric factorizations, respectively, in their rows. Since 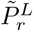 and 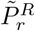 are non-negative matrices, we require the matrices 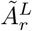, 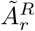, 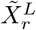, and 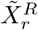 to be non-negative as well. The factorizations were computed by using the Matlab N-way Toolbox [6].

### 4.8 Inter-hemispheric functional connectivity

To assess the inter-hemispheric functional organization of the identified generators, we proceed as follows. Note first that the generator time-courses 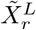 and 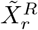 of the left- and right-hemispheres of run *r* can be written as

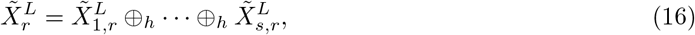

and

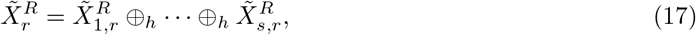

where 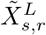, and 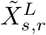, contain the left- and right-hemispheric generator time-courses of subject *s*. For a given left-hemispheric generator, and for each subject *s* and run *r*, we calculate the Pearson correlation coefficients between its time-course, which corresponds to one of the rows of 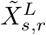, and the power fluctuations of all right-hemispheric cortical vertices, which correspond to the columns of 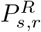. This yields an *n*/2-dimensional column vector 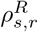 of inter-hemispheric correlation coefficients. By repeating the same calculation for the corresponding generator in the right hemisphere, we obtain a similar correlation vector 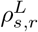, which can subsequently be combined into the n-dimensional correlation vector

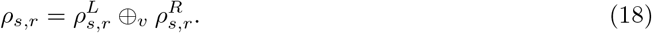

Thus, the *v*-th entry of *ρ_s,r_* contains the Pearson correlation coefficient between the power-fluctuations of subject *s* and run *r* at vertex *v* and the contra-lateral time-course of the selected generator. Run-specific population-level correlation vectors *ρ_r_* are now obtained by averaging the Fisher-transformed subject-specific correlation vectors over subjects:

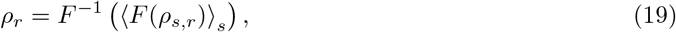

where *F* and *F*^−1^ denote the Fisher and inverse Fisher transformations, respectively, and <>_*s*_ denote averaging over s (subjects). Although in correlating the generator time-courses with the low-frequency power fluctuations, the (absolute) power matrices *P_r_* are used, the generator time-courses themselves were obtained by factorizing the power-density matrices 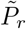, which leads to uninformative negative correlations in *ρ_r_*. Therefore, the negative entries in *ρ_r_* were set to zero, yielding rectified correlation vectors 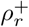.

## Acknowledgments

RH and GD were funded by the European Research Council (Advanced Grant DYSTRUCTURE No.295129), the Spanish Research Project PSI2013-42091-P, the CONSOLIDER-INGENIO 2010 Program CSD2007-00012, and the FP7-ICT Brainscales (269921) awared to GD. DM was funded by the Wellcome Trust (101253/Z/13/Z), the KU Leuven Special Research Fund (C16/15/070), and the Research Foundation Flanders (FWO) (GOF76.16N and G0936.16N). Data were provided by the Human Connectome Project, WU-Minn Consortium (Principal Investigators: David Van Essen and Kamil Ugurbil; 1U54MH091657) funded by the 16 NIH Institutes and Centers that support the NIH Blueprint for Neuroscience Research; and by the McDonnell Center for Systems Neuroscience at Washington University.

## References

[1] L. I. Aftanas and S. A. Golocheikine. Human anterior and frontal midline theta and lower alpha reflect emotionally positive state and internalized attention: High-resolution EEG investigation of meditation. Neurosci. Lett., 310(1):57–60, 2001.

[2] J. Ahveninen. The long reach of the gene. Psychologist, 26(3):194–198, 2013.

[3] A. P. Baker, M. J. Brookes, I. A. Rezek, S. M. Smith, T. Behrens, P. J. P. Smith, and M. Woolrich. Fast transient networks in spontaneous human brain activity. Elife, 2014(3):1–18, 2014.

[4] M. C. M. Bastiaansen, K. B. E. Bocker, C. H. M. Brunia, J. C. De Munck, and H. Spekreijse. Event-related desynchronization during anticipatory attention for an upcoming stimulus: A comparative EEG/MEG study. Clin. Neurophysiol., 112(2):393–403, 2001.

[5] H. Berger. Uber das elektrenkephalogramm des menschen. 278(1875), 1929.

[6] R. Bro. PARAFAC. Tutorial and applications. Chemom. Intell. Lab. Syst., 38(2):149–171, 1997.

[7] Matthew J Brookes, Mark Woolrich, Henry Luckhoo, Darren Price, Joanne R Hale, Mary C Stephenson, Gareth R Barnes, Stephen M Smith, and Peter G Morris. Investigating the electrophysiological basis of resting state networks using magnetoencephalography. Proc. Natl. Acad. Sci. U. S. A., 108(40):16783–8, oct 2011.

[8] F. Brunet, A. Bartoli, and L. Uda. L-tangent norm: A low computational cost criterion for choosing regularization weights and its use for range surface reconstruction. Proc. Fourth Int. Symp. 3D Date Process. Vis. Transm., 2008.

[9] A. Capilla, J. M. Schoffelen, G. Paterson, G. Thut, and J. Gross. Dissociated alpha-band modulations in the dorsal and ventral visual pathways in visuospatial attention and perception. Cereb. Cortex, 24(2):550–561, 2014.

[10] C. Ciulla, T. Takeda, and H. Endo. MEG Characterization of Spontaneous Alpha Rhythm in the Human Brain MEG Characterization of Spontaneous Alpha Rhythm in the Human Brain. (March), 1999.

[11] W. A. Cobb, R. J. Guiloff, and J. Cast. Breach rhythm: The EEG related to skull defects. Electroencephalogr. Clin. Neurophysiol., 47(3):251–271, 1979.

[12] D. Cohen. Magnetoencephalography: evidence of magnetic fields produced by alpha rhythm currents, 1968.

[13] G. L. Colclough, M. J. Brookes, S. M. Smith, and M. W. Woolrich. A symmetric multivariate leakage correction for MEG connectomes. Neuroimage, 117:439–448, 2015.

[14] M. Corbetta and G. L. Shulman. Control of Goal-Directed and Stimulus-Driven Attention in the Brain. Nat. Rev. Neurosci., 3(3):215–229, 2002.

[15] S. Della Penna, K. Torquati, V. Pizzella, C. Babiloni, R. Franciotti, P. M. Rossini, and G. L. Romani. Temporal dynamics of alpha and beta rhythms in human SI and SII after galvanic median nerve stimulation. A MEG study. Neuroimage, 22(4):1438–1446, 2004.

[16] B. A. Diaz, S. Van Der Sluis, S. Moens, J. S. Benjamins, F. Migliorati, D. Stoffers, A. Den Braber, S.-S. Poil, R. Hardstone, D. Van’t Ent, D. I. Boomsma, E. De Geus, H. D. Mansvelder, E. J. W. Van Someren, and K. Linkenkaer-Hansen. The Amsterdam Resting-State Questionnaire reveals multiple phenotypes of resting-state cognition. Front. Hum. Neurosci., 7(August):446, 2013.

[17] L. A. Finelli, P. Ackermann, and A. A. Borbely. Individual ‘fingerprints’ in human sleep EEG topography. Neuropsychopharmacology, 25(01):S57–S62, 2001.

[18] E. S. Finn, X. Shen, D. Scheinost, M. D. Rosenberg, J. Huang, M. M. Chun, X. Papademetris, and R. T. Constable. Functional connectome fingerprinting: identifying individuals using patterns of brain connectivity. Nat. Neurosci., 18(11):1664–1671, oct 2015.

[19] J. N. Frey, N. Mainy, J.-P. Lachaux, N. Müller, O. Bertrand, and N. Weisz. Selective modulation of auditory cortical alpha activity in an audiovisual spatial attention task. J. Neurosci., 34(19):6634–9, 2014.

[20] Matthew F. Glasser, Stamatios N. Sotiropoulos, J. Anthony Wilson, Timothy S. Coalson, Bruce Fischl, Jesper L. Andersson, Junqian Xu, Saad Jbabdi, Matthew Webster, Jonathan R. Polimeni, David C. Van Essen, and Mark Jenkinson. The minimal preprocessing pipelines for the Human Connectome Project. Neuroimage, 80:105–124, 2013.

[21] R. I. Goldman, J. M. Stern, J. Engel, and M. S. Cohen. Simultaneous EEG and fMRI of the alpha rhythm. Neuroreport, 13(18):2487–92, 2002.

[22] C. M. Gómez, J. Marco-Pallarés, and C. Grau. Location of brain rhythms and their modulation by preparatory attention estimated by current density. Brain Res., 1107(1):151–160, 2006.

[23] S. I. Gonéalves, J. C. De Munck, P. J. W. Pouwels, R. Schoonhoven, J. P. A. Kuijer, N. M. Maurits, J. M. Hoogduin, E. J. W. Van Someren, R. M. Heethaar, and F. H. Lopes Da Silva. Correlating the alpha rhythm to BOLD using simultaneous EEG/fMRI: Inter-subject variability. Neuroimage, 30(1):203–213, 2006.

[24] S. Haegens, B. F. Handel, and O. Jensen. Top-Down Controlled Alpha Band Activity in Somatosensory Areas Determines Behavioral Performance in a Discrimination Task. J. Neurosci., 31(14):5197–5204, 2011.

[25] E. L. Hall, M. W. Woolrich, C. E. Thomaz, P. G. Morris, and M. J. Brookes. Using variance information in magnetoencephalography measures of functional connectivity. Neuroimage, 67:203–212, 2013.

[26] P.C. Hansen and D.P. O’Leary. The use of the L-curve in the regularization of discrete ill-posed problems. SIAM J. Sci. Comput., 14(6):1487–1503, 1993.

[27] R. Hari and R. Salmelin. Human cortical oscillation: a neuromagnetic view through the skull. Trends Neurosci., 20(1):44–49, 1997.

[28] B. Herrmann, M. J. Henry, S. Haegens, and J. Obleser. Temporal expectations and neural amplitude fluctuations in auditory cortex interactively influence perception. Neuroimage, 124:487–497, 2016.

[29] R. Hindriks, X.D. Arsiwalla, T. Panagiotaropoulos, M. Besserve, P.F.M.J. Verschure, N.K. Logothetis, and G. Deco. Discrepancies between Multi-Electrode LFP and CSD Phase-Patterns: A Forward Modeling Study. Front. Neural Circuits, 10(July):51, 2016.

[30] J.F. Hipp, D.J. Hawellek, M. Corbetta, M. Siegel, and A.K. Engel. Large-scale cortical correlation structure of spontaneous oscillatory activity. Nat. Neurosci., 15(6):884–90, jun 2012.

[31] M. X. Huang, C. W. Huang, A. Robb, A. M. Angeles, S. L. Nichols, D. G. Baker, T. Song, D. L. Harrington, R. J. Theilmann, R. Srinivasan, D. Heister, M. Diwakar, J. M. Canive, J. C. Edgar, Y. H. Chen, Z. Ji, M. Shen, F. El-Gabalawy, M. Levy, R. McLay, J. Webb-Murphy, T. T. Liu, A. Drake, and R. R. Lee. MEG source imaging method using fast L1 minimum-norm and its applications to signals with brain noise and human resting-state source amplitude images. Neuroimage, 84:585–604, 2014.

[32] R. Ishii, K. Shinosaki, S. Ukai, T. Inouye, T. Ishihara, T. Yoshimine, N. Hirabuki, H. Asada, T. Kihara, S. E. Robinson, and M. Takeda. Medial prefrontal cortex generates frontal midline theta rhythm. Neuroreport, 10(4):675–679, 1999.

[33] O. Jensen, M. Bonnefond, and R. VanRullen. An oscillatory mechanism for prioritizing salient unattended stimuli. Trends Cogn. Sci., 16(4):200–6, apr 2012.

[34] O. Jensen and A. Mazaheri. Shaping functional architecture by oscillatory alpha activity: gating by inhibition. Front. Hum. Neurosci., 4(November):186, jan 2010.

[35] S. P. Kelly, E. C. Lalor, R. B. Reilly, and J. J. Foxe. Increases in alpha oscillatory power reflect an active retinotopic mechanism for distracter suppression during sustained visuospatial attention. J. Neurophysiol., 95(6):3844–51, jun 2006.

[36] W. Klimesch, P. Sauseng, and S. Hanslmayr. EEG alpha oscillations: the inhibition-timing hypothesis. Brain Res. Rev., 53(1):63–88, jan 2007.

[37] G. G. Knyazev, A. N. Savostyanov, A. V. Bocharov, S. S. Tamozhnikov, and A. E. Saprigyn. Task-positive and task-negative networks and their relation to depression: EEG beamformer analysis, volume 306. Elsevier B.V., 2016.

[38] A. Kondacs and M. Szabo. Long-term intra-individual variability of the background EEG in normals. Clin. Neurophysiol., 110(10):1708–1716, 1999.

[39] L. J. Larson-Prior, R. Oostenveld, S. Della Penna, G. Michalareas, F. Prior, A. Babajani-Feremi, J. M. Schoffelen, L. Marzetti, F. de Pasquale, F. Di Pompeo, J. Stout, M. Woolrich, Q. Luo, R. Bucholz, P. Fries, V. Pizzella, G. L. Romani, M. Corbetta, and A. Z. Snyder. Adding dynamics to the Human Connectome Project with MEG. Neuroimage, 80:190–201, 2013.

[40] H. Laufs, A. Kleinschmidt, A. Beyerle, E. Eger, A. Salek-Haddadi, C. Preibisch, and K. Krakow. EEG-correlated fMRI of human alpha activity. Neuroimage, 19:1463–1476, aug 2003.

[41] L. Lehtela, R. Salmelin, and R. Hari. Evidence for reactive magnetic 10-Hz rhythm in the human auditory cortex. 222:111–114, 1997.

[42] M. Liljestrom, J. Kujala, O. Jensen, and R. Salmelin. Neuromagnetic localization of rhythmic activity in the human brain: A comparison of three methods. Neuroimage, 25(3):734–745, 2005.

[43] K. Linkenkaer-Hansen, V. V. Nikouline, J. M. Palva, and R. J. Ilmoniemi. Long-range temporal correlations and scaling behavior in human brain oscillations. J. Neurosci., 21(4):1370–7, 2001.

[44] Q. Liu, S. Farahibozorg, and C. Porcaro. Detecting large-scale networks in the human brain using high-density electroencephalography. BioRxiv, pages 1–31, 2016.

[45] S. Makeig, M. Westerfield, T. P. Jung, S. Enghoff, J. Townsend, E. Courchesne, and T. J. Sejnowski. Dynamic brain sources of visual evoked responses. Science (80-.)., 295(5555):690–694, 2002.

[46] I. Manshanden, J. C. De Munck, N. R. Simon, and F. H. Lopes da Silva. Source localization of MEG sleep spindles and the relation to sources of alpha band rhythms. Clin. Neurophysiol., 113(12):1937–1947, 2002.

[47] D. Mantini, S. D. Penna, L. Marzetti, F. de Pasquale, V. Pizzella, M. Corbetta, and G. L. Romani. A Signal-Processing Pipeline for Magnetoencephalography Resting-State Networks. Brain Connect., 1(1):49–59, 2011.

[48] D. Mantini, M.G. Perrucci, C. Del Gratta, G.L. Romani, and M. Corbetta. Electrophysiological signatures of resting state networks in the human brain. Proc. Natl. Acad. Sci., 104(32):13170, 2007.

[49] S. D. Mayhew, D. Ostwald, C. Porcaro, and A. P. Bagshaw. Spontaneous EEG alpha oscillation interacts with positive and negative BOLD responses in the visual-auditory cortices and defaultmode network. Neuroimage, 76:362–372, 2013.

[50] A. Mazaheri, M. R. van Schouwenburg, A. Dimitrijevic, D. Denys, R. Cools, and O. Jensen. Region-specific modulations in oscillatory alpha activity serve to facilitate processing in the visual and auditory modalities. Neuroimage, 87:356–362, 2014.

[51] F. Miwakeichi, E. Mart?nez-Montes, P. A. Valdes-Sosa, N. Nishiyama, H. Mizuhara, and Y. Yamaguchi. Decomposing EEG data into space-time-frequency components using Parallel Factor Analysis. Neuroimage, 22(3):1035–45, jul 2004.

[52] J. Mo, Y. Liu, H. Huang, and M. Ding. Coupling between visual alpha oscillations and default mode activity. Neuroimage, 68:112–118, 2013.

[53] T. Montez, S.-S. Poil, B. F. Jones, I. Manshanden, J. P. A. Verbunt, B. W. van Dijk, A. B. Brussaard, A. van Ooyen, C. J. Stam, P. Scheltens, and K. Linkenkaer-Hansen. Altered temporal correlations in parietal alpha and prefrontal theta oscillations in early-stage Alzheimer disease. Proc. Natl. Acad. Sci. U. S. A., 106(5):1614–1619, 2009.

[54] N. Muller and N. Weisz. Lateralized auditory cortical alpha band activity and interregional connectivity pattern reflect anticipation of target sounds. Cereb. Cortex, 22(7):1604–1613, 2012.

[55] M. Näpflin, M. Wildi, and J. Sarnthein. Test-retest reliability of resting EEG spectra validates a statistical signature of persons. Clin. Neurophysiol., 118(11):2519–2524, 2007.

[56] L. Narici, N. Forss, V. Jousmaki, M. Peresson, and R. Hari. Evidence for a 7- to 9-Hz Sigma Rhythm in the Human SII Cortex. 668:662–668, 2001.

[57] E. Niedermeyer. Alpha-like rhythmical activity of the temporal lobe. Clin Electroencephalogr, 21(4):210–224, 1990.

[58] R. Oostenveld, P. Fries, E. Maris, and J.-M. Schoffelen. FieldTrip: Open source software for advanced analysis of MEG, EEG, and invasive electrophysiological data. Comput. Intell. Neurosci., page 156869, jan 2011.

[59] Y. Petrov. Harmony: EEG/MEG linear inverse source reconstruction in the anatomical basis of spherical harmonics. PLoS One, 7(10):e44439, jan 2012.

[60] M. Ploner, J. Gross, L. Timmermann, B. Pollok, and A. Schnitzler. Pain suppresses spontaneous brain rhythms. Cereb. Cortex, 16(4):537–540, 2006.

[61] V. Romei, V. Brodbeck, C. Michel, A. Amedi, A. Pascual-Leone, and G. Thut. Spontaneous fluctuations in posterior alpha-band EEG activity reflect variability in excitability of human visual areas. Cereb. cortex, 18(9):2010–8, sep 2008.

[62] S. Salenius, M. Kajola, W. L. Thompson, S. Kosslyn, and R. Hari. Reactivity of magnetic parieto-occipital alpha rhythm during visual imagery. Electroencephalogr. Clin. Neurophysiol., 95(6):453–462, 1995.

[63] S. Salenius, A. Schnitzler, R. Salmelin, V. Jousmäki, and R. Hari. Modulation of human cortical rolandic rhythms during natural sensorimotor tasks. Neuroimage, 5(3):221–228, 1997.

[64] R. Salmelin and R. Hari. Spatiotemporal characteristics of sensorymotor neuromagnetic rhythms related to thumb movement. Neuroscience, 60(2):537–550, 1994.

[65] R. Scheeringa, M. C. M. Bastiaansen, K. M. Petersson, R. Oostenveld, D. G. Norris, and P. Hagoort. Frontal theta EEG activity correlates negatively with the default mode network in resting state. Int. J. Psychophysiol., 67(3):242–251, 2008.

[66] A. C. Snyder and J. J. Foxe. Anticipatory Attentional Suppression of Visual Features Indexed by Oscillatory Alpha-Band Power Increases: A High-Density Electrical Mapping Study. J. Neurosci., 30(11):4024–32, 2010.

[67] V. Stormer. Salient, irrelevant sounds reflexively induce alpha rhythm desynchronization in parallel with slow potential shifts in visual cortex. J. Cogn. Neurosci., 26(3):194–198, 2016.

[68] J. Tiihonen, R. Hari, M. Kajola, J. Karhu, S. Ahlfors, and S. Tissari. Magnetoencephalographic 10-Hz rhythm from the human auditory cortex. Neurosci. Lett., 129(2):303–305, 1991.

[69] D. C. Van Essen, S. M. Smith, D. M. Barch, T. E. J. Behrens, E. Yacoub, and K. Ugurbil. The WU-Minn Human Connectome Project: An overview. Neuroimage, 80:62–79, 2013.

[70] N. Weisz, T. Hartmann, N. Müller, I. Lorenz, and J. Obleser. Alpha rhythms in audition: Cognitive and clinical perspectives. Front. Psychol., 2(APR):1–15, 2011.

[71] Nathan Weisz, Nadia M??ller, Sabine Jatzev, and Olivier Bertrand. Oscillatory alpha modulations in right auditory regions reflect the validity of acoustic cues in an auditory spatial attention task. Cereb. Cortex, 24(10):2579–2590, 2014.

[72] S.J. Williamson, L. Kaufman, Z.L. Lu, J.Z. Wang, and D. Karron. Study of human occipital alpha rhythm: the alphon hypothesis and alpha suppression. Int. J. Psychophysiol., 26:63–76, 1997.

[73] M. Wöstmann, B. Herrmann, B. Maess, and J. Obleser. Spatiotemporal dynamics of auditory attention synchronize with speech. Proc. Natl. Acad. Sci. U. S. A., 113(14):1523357113-, 2016.

[74] N. Yamagishi, N. Goda, D. E. Callan, S. J. Anderson, and M. Kawato. Attentional shifts towards an expected visual target alter the level of alpha-band oscillatory activity in the human calcarine cortex. Cogn. Brain Res., 25(3):799–809, 2005.

